# Compensatory evolution facilitates the acquisition of multiple plasmids in bacteria

**DOI:** 10.1101/187070

**Authors:** Alfonso Santos-Lopez, Cristina Bernabe-Balas, Alvaro San Millan, Rafael Ortega-Huedo, Andreas Hoefer, Manuel Ares-Arroyo, Bruno Gonzalez-Zorn

**Author notes:** Current address: Department of Microbiology and Molecular Genetics, University of Pittsburgh, Pittsburgh, Pennsylvania, USA. Current address: Department of Microbiology, Ramón y Cajal University Hospital (IRYCIS), and CIBERESP. 28034, Madrid, Spain (Alvaro San Millan).

## Abstract

The coexistence of multicopy plasmids is a common phenomenon. However, the evolutionary forces promoting these genotypes are poorly understood. In this study, we have analyzed multiple ColE1 plasmids (pB1000, pB1005 and pB1006) coexisting within *Haemophilus influenzae* RdKW20 in all possible combinations. When transformed into the naïve host, each plasmid type presented a particular copy number and produced a specific resistance profile and biological cost, whether alone or coexisting with the other plasmids. Therefore, there was no fitness advantage associated with plasmid coexistence that could explain these common plasmid associations in nature. Using experimental evolution, we showed how *H. influenzae* Rd was able to completely compensate the fitness cost produced by any of these plasmids. Crucially, once the bacterium has compensated for a first plasmid, the acquisition of new multicopy plasmid(s) did not produced any extra biological cost. We argue therefore that compensatory adaptation pave the way for the acquisition of multiple coexisting ColE1 plasmids.

**Importance:** Antibiotic resistance is a major concern for human and animal health. Plasmids play a major role in the acquisition and dissemination of antimicrobial resistance genes. In this report we investigate, for the first time, how plasmids are capable to cohabit stably in populations. This coexistence of plasmids is driven by compensatory evolution alleviating the cost of a first plasmid, which potentiates the acquisition of further plasmids at no extra cost. This phenomenon explains the high prevalence of plasmids coexistance in wild type bacteria, which generates multiresistant clones and contributes to the maintenance and spread of antibiotic resistance genes within bacterial populations.

## Introduction

Antibiotic resistance is a serious problem in animal and human health and bacterial plasmids play an essential role in the dissemination of resistance (1). In last years, numerous works have described the importance of small ColE1-like plasmids in the dissemination of resistance genes (2-15). These plasmids replicate via two RNAs (16). Natural SNPs in these RNAs allow different ColE1-like plasmids to stably cohabit within the same cell (17, 18). If the plasmids bear antibiotic resistance genes, this cohabitation confers antibiotic multiresistance to the host bacteria (9).

The acquisition of plasmids usually entails a biological cost to the host bacterium that will generate a selection against plasmid-bearing clones (19, 20). Thus, it is reasonable to assume that the accumulation of various plasmids will decrease the fitness of bacteria and therefore clones bearing several plasmids will be outcompeted in bacterial populations. Notwithstanding, ColE1 plasmids cohabitation is a common phenomenon in nature (9, 11, 13, 14, 21). In this study we test two hypotheses that could explain the high prevalence of plasmid coexistence in nature: i) Positive epistasis: the cost imposed by multiple ColE1 plasmids is lower than the addition of the costs produced by each of the plasmids alone (22) and ii) compensatory evolution increasing permissiveness to ColE1-like plasmids: once the bacterium has compensated the cost of a single plasmid the acquisition of new replicons does not affect the bacterial fitness (23).

Here, we demonstrated that once the fitness cost of a ColE1-like plasmid is compensated the acquisition of more ColE1-like plasmids does not incur any biological cost to the bacteria, facilitating plasmid cohabitation.

## Materials and Methods

### Bacterial strains, culture conditions and antibiotic susceptibility determination

All strains and plasmids used in this study are listed in Table S1. *H. influenzae* was electroporated with pB1000, pB1005 and/or pB1006 from *P. multocida* BB1044 (9) as previously described (10). *H. influenzae* was cultured on chocolate agar PolyViteX plates (BioMérieux, France) and in *Haemophilus* Test Medium (HTM) broth (Wider, Francisco Soria Melguizo, Spain) shaking at 125 RPM and 37° C in microaerophilic conditions (5% CO_2_). Antibiotic susceptibility was determined via minimal inhibitory concentration (MIC) of ampicillin, streptomycin and tetracycline by broth microdilution according to the CLSI guidelines (24).

### Plasmid stability

We assessed the stability of the plasmids in all ColE1-bearing strains (Table 1). All three plasmids in the seven combinations presented 100% stability in *H. influenzae* Rd after 200 generations.

**Table 1.** Plasmid copy number, relative fitness and MIC of the transformed plasmids in Rd.

| Strain | Plasmid Copy Number | Relative Fitness<sup>a</sup> | MIC(mg/l)<sup>b</sup> |  |  |
| --- | --- | --- | --- | --- | --- |
|  |  |  | Tet | Str | Amp |
| Rd | 0 | 1 | 2 | 8 | 0,25 |
| Rd/pB1006 | 10,53 ± 1,112 | 0,979 ± 0,012 | 16 | 8 | 0,25 |
| Rd/pB1005 | 20,45 ± 2,59 | 0,950 ± 0,013 | 2 | 1024 | 0,25 |
| Rd/pB1000 | 25,02 ± 1,92 | 0,946 ± 0,002 | 2 | 8 | 512 |
| Rd/pB1005/pB1006 | 28,17 ± 2,37 | 0,921 ± 0,017 | 16 | 1024 | 0,25 |
| Rd/pB1006/pB1000 | 33,96 ± 3,01 | 0,909 ± 0,003 | 16 | 8 | 512 |
| Rd/pB1005/pB1000 | 43,22 ± 4,43 | 0,899 ± 0,009 | 2 | 512 | 512 |
| Rd/pB1006/pB1005/pB1006 | 53,58 ± 12,48 | 0,883 ± 0,016 | 32 | 512 | 512 |

Plasmid copy number, fitness and antimicrobial susceptibility of the seven strains generated eletroporing the plasmids in the Rd strain. ^a^ The relative fitness is expressed compared to the Rd strain. ^b^ Tet, tetracycline; Str, streptomycin and Amp, ampicillin.

### Plasmid copy number quantification

The average plasmid copy number per cell was determined by quantitative PCR (qPCR) as described by San Millan *et al.* (25). To determine the plasmid copy number five independent DNA extractions were performed for each strain and qPCR was then carried out in triplicate for each extraction. Each strain was grown in 2 ml of fresh HTM and the DNA was extracted at an OD600 of approximately 0.9 using the QIAamp DNA Mini Kit (Qiagen, Inc, Chatworth, California, USA). The DNA was quantified using a Nanodrop. Following the indications of Providenti *et al.* (26), digested DNA is a better template for plasmid quantification by qPCR than non-digested DNA. Therefore, plasmids were linearized with *PstI* (Takara, Japan) for 2 hours at 37°C. In order to determine the average plasmid copy number per chromosome, the chromosomal monocopy gene *rpoB* was amplified to compare the ratio of plasmid-chromosomal DNA.

qPCRs were performed using a My iQ Single Color Real-Time PCR Detetion System (Bio-Rad laboratories) with the iQ SYBR Green Supermix (Bio-Rad Laboratories) at a final DNA concentration of 10 pg/μl. The reaction efficiency was calculated for each reaction based on the standard curve generated by performing a qPCR with five 8-fold dilutions of the template DNA in triplicate (~0.2 ng/μl to 50 fg/μl working range of DNA concentration), and reactions with an R^2^ lower than 0.985 were discarded. All primers used in the reactions, as well as the melting temperatures and efficiencies of the qPCRs are described in Table S2. The amplification conditions were as follows: initial denaturation for 10 min at 94°C, followed by 30 cycles of denaturation for 1’ at 94°C, annealing for 1’ at 51,7°C (*rpoB*) or 58,5°C (pB1000) 55,1°C (pB1005) and 54,6°C (pB1006) and extension for 1’ at 72°C. Inter-run calibration samples were used to normalize the results from different plates of each qPCR. To calculate plasmid copy number per chromosome we used the following formula:

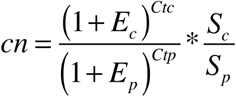

Where *cn* is the plasmid copy number per chromosome, *S*_*c*_ and *S*_*p*_ are the sizes of the chromosomal and plasmid amplicons (in bp), *E*_*c*_ and *E*_*p*_ are the efficiencies of the chromosomal and plasmid qPCRs (relative to 1), and *Ctc* and *Ctp* are the threshold cycles of the chromosomal and plasmid reactions, respectively.

### Fitness determination

Bacterial fitness was determined by direct competition experiments between *H. influenzae* Rd and *H. influenzae* Rd bearing plasmid(s) in HTM medium in 3-5 independent experiments (25). Strains were grown for 16 hours at 37°C and 5% CO_2_ in HTM, then 10^6^ CFU of each competitor were suspended in 2 ml of HTM broth. The inocula was grown at 37° C, 5% CO_2_ and 125 RPM for 24 hours, after which 10^6^ CFU were transferred to 2 ml of fresh HTM every 24 hours (1/1000) for 5 days, resulting in 10 generations per serial passage. Samples were taken at time 0 and every 24 hours for 5 days. Aliquots were then plated onto non-selective chocolate agar, and the ratio of the two competing strains was measured by replica plating 50-100 colonies on chocolate agar plates containing ampicillin, streptomycin and/or tetracycline corresponding the resistance gene(s) borne by the plasmid(s). The selection coefficients (*s*) were calculated using the regression model *s* = ln(CI/t), where the CI (competition index) was calculated everyday as the ratio between the CFU of the resistant and susceptible strains at *t1* divided by the same ratio at *t0*. Time (t) was calculated as the log2 of the dilution factor (i.e. number of bacterial generations). Relative fitness (W) was calculated as 1-*s*.

Epistasis among plasmids was determined as described by Hall *et al.* (27). Epistasis (*ɛ*) can be calculated as *ɛ* = W_(plasmid A;plasmid B)_-W_(plasmid A)_ × W_(plasmid B)_, where W_(plasmid A)_ or W_(plasmid B)_ is the relative fitness of the strain bearing plasmid 1 or plasmid 2 compared to the plasmid free strain and W(plasmid A;plasmid B) is the relative fitness of the strain bearing both plasmids relative to the same plasmid free strain. Then, the propagation error (σ_*ɛ*_) was calculated by applying with the formula:

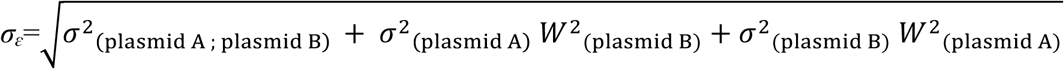

If the epistasis value is within the propagation error range, there are no significant epistatic interactions between the plasmids. If on the other hand the value is beyond the range of the propagation error one can assume that there are epistatic interactions. Both formulas were adjusted to estimate the epistasis among the three plasmids.

### *In silico* analysis

We analyzed all Pasteurellaceae and Enterobacteriaceae genomes available in the GenBank database as of November 2015. Only genomes with the status “Complete” were included in the analysis.

## Results and discussion

### Coexistence of multicopy plasmids is common in nature

ColE1 plasmids are found mainly in the families of bacteria Enterobacteriaceae and Pasteurellaceae (2, 4, 7, 9, 15, 28-30). We combined *in silico* and experimental information to analyze the prevalence of ColE1 plasmid coexistence in nature.

We scanned the presence of ColE1 plasmids in the Enterobacteriaceae family where they were first described (16). We detected 631 plasmids in databases, in 490 complete genomes. 100 plasmids (16% of the total) belonged to the ColE1 superfamily. 45% of ColE1 plasmids coexisted with at least another ColE1 plasmid in the same cell (Table S1). We analyzed the distribution of ColE1 plasmids described in the Enterobacteriaceae genomes. As described by San Millan *et al.* (22) if plasmids were distributed randomly, they would follow a Poisson distribution across bacterial hosts. If there is any factor influencing the plasmid distribution such as conjugation, epistasis or selection, the observed ratio of plasmid per genome may suffer a significant deviation from the expected Poisson distribution. We found significantly different pattern between the observed and the expected distribution of genomes bearing zero, one, and two or more ColE1 plasmids: (chi-square test, P < 0.001, χ^2^ = 0.903, df = 2) (Figure 1). Observed strains lacking plasmids are as common as expected, while strains carrying only one ColE1 replicon were underrepresented and the strains bearing 2 or more ColE1 plasmids were overrepresented. These results confirm a tendency towards ColE1 plasmid coexistence in enterobacteria.

**Figure 1.**
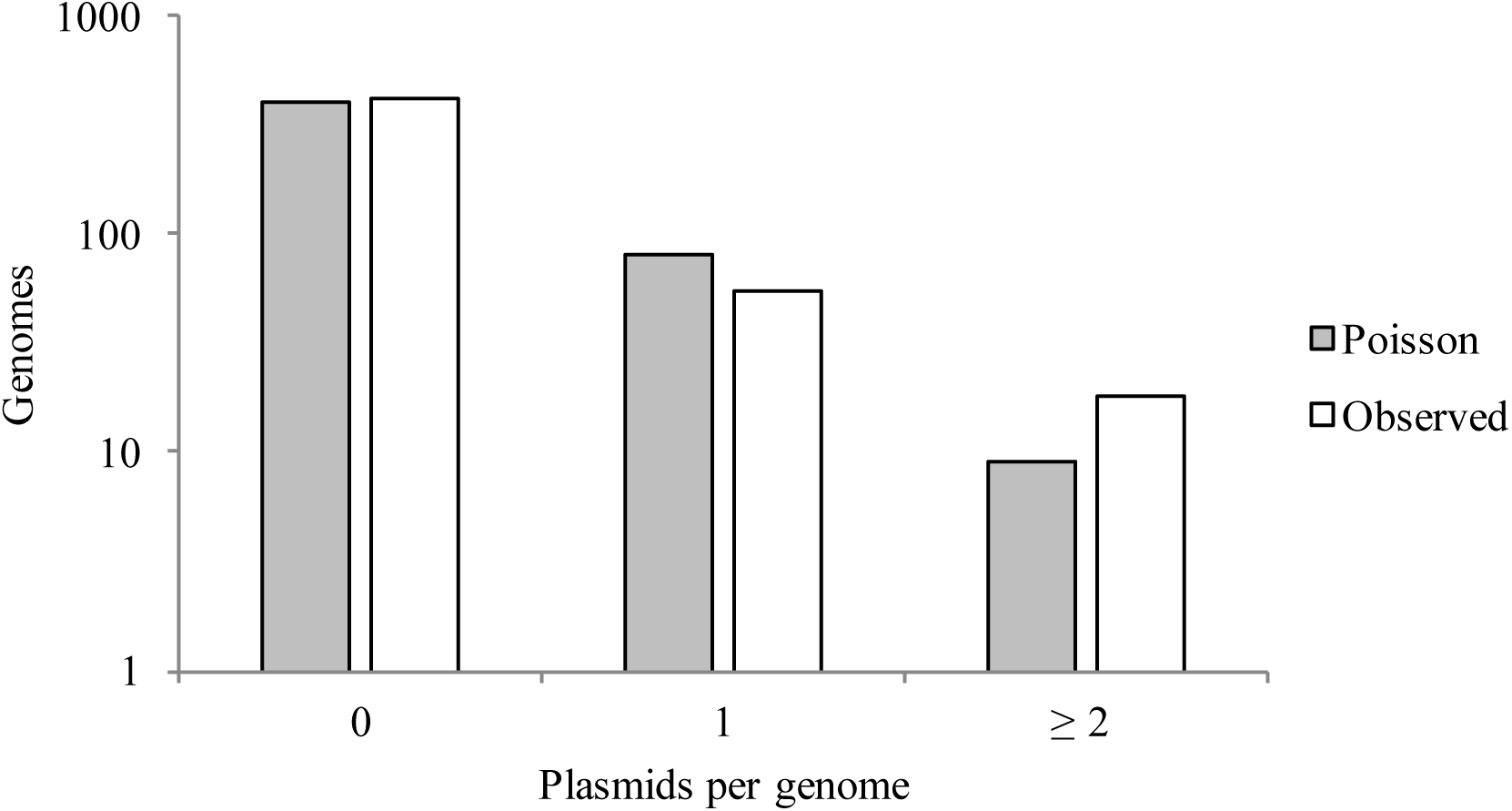
Distribution of ColE1 plasmids in bacterial genomes. Number of plasmids per bacterial genome. The grey bars represent the expected number of genomes carrying from zero to ≥2 plasmids following a Poisson distribution (using the average plasmid/strain observed in the 490 bacterial genomes analyzed from GenBank). The white bars represent the observed frequency of strains carrying zero to ≥2 plasmids in the bacterial genomes analyzed.

In Pasteurellaceae there were too few genomes available in databases for a robust analysis. However, previous works suggested that ColE1 plasmid coexistence is also common (9-12, 31, 32). Actually, plasmid pB1000, which is the most widely distributed ColE1-like plasmid in this family, has been always described coexisting with other ColE1 plasmids: in *Haemophilus parasuis* (32), *Pasteurella aerogenes* (33) *Actinobacillus pleuropneumoniae* (Accesion number to GenBank: CP001904) and *Pasteurella multocida* (9) (see Table S1). Taken together, our data showed that coexistence of ColE1-like plasmids is a frequent event in nature.

### Biological cost of ColE1 plasmids has a multiplicative effect

Plasmids produce a fitness cost in bacteria (19, 20). It is reasonable to assume that bacteria bearing two (or three) plasmids would present a decrease in fitness compared to bacteria bearing only one plasmid. However, Silva *et al.* (34) and San Millan *et al.* (22) have demonstrated that, in some cases, the presence of one plasmid favors the presence of a second: once a bacterium had acquired a plasmid, the presence of a second (or third) plasmid in the host does not incurred an additional significant biological cost, even when these plasmids produced a cost when they are alone in the cell. This phenomenon is known as positive epistasis between coexisting plasmids.

To determine whether epistatic interactions can explain ColE1 coexistence, we transformed *H. influenzae* RdKw20 (Rd hereinafter) by electroporation with the plasmids pB1000, pB1005 and pB1006, recovered from the a clinical isolate of *P. multocida* BB1044 (9). Thus, we produced a model of bacteria with seven possible plasmid combinations (one, two or three different ColE1 plasmids per bacterium). All three plasmids are composed by a variable region in which antibiotic resistance genes are encoded -*bla*_ROB-1_ in pB1000, *str*A in pB1005 and the *tet*(O) in pB1006-and a highly similar conserved region with the plasmid housekeeping functions (Figure S1).

To estimate the biological cost associated with the replicon(s), we performed direct competition assays between *H. influenzae* Rd strain bearing the plasmid(s) and the plasmid-free *H. influenzae* Rd strain in culture medium lacking antibiotic pressure. As expected, bacteria bearing one or more plasmid(s) were less fit than the ancestral strain (Table 1 and Figure 2). In order to test if epistatic interactions were able to explain coexistence of ColE1 replicons, we calculated epistasis as described by Hall *et al.* (27) (see methods). No significant epistatic interactions were found across the four combinations: pB1005/pB1006 (*ɛ* = −0.009, σ_ɛ_ = ± 0.0425), pB1000/pB1006 (*ɛ* = −0.017, σ_ɛ_ = ± 0.042), pB1000/pB1005 (*ɛ* = 0.000, σ_ɛ_ = ± 0.025) and pB1000/pB1005/pB1006 (*ɛ* = 0.0002, σ_ɛ_ = ± 0.021). (Figure 2). The biological cost of ColE1 plasmids produced a multiplicative effect: the relative fitness of strains carrying two or more ColE1 plasmids (e.g. W_(pB1000, pB10056)_ = 0.899) was not significantly different form a multiplication of the relative fitnesses of the two bacteria bearing each one of the plasmids (e.g. W_(pB1000)_ × W_(pB1006)_ = 0.946 × 0.979 = 0.899). These results therefore suggest that epistatic interactions among recently acquired plasmids are may not be at the origin of ColE1 plasmid coexistence.

**Figure 2.**
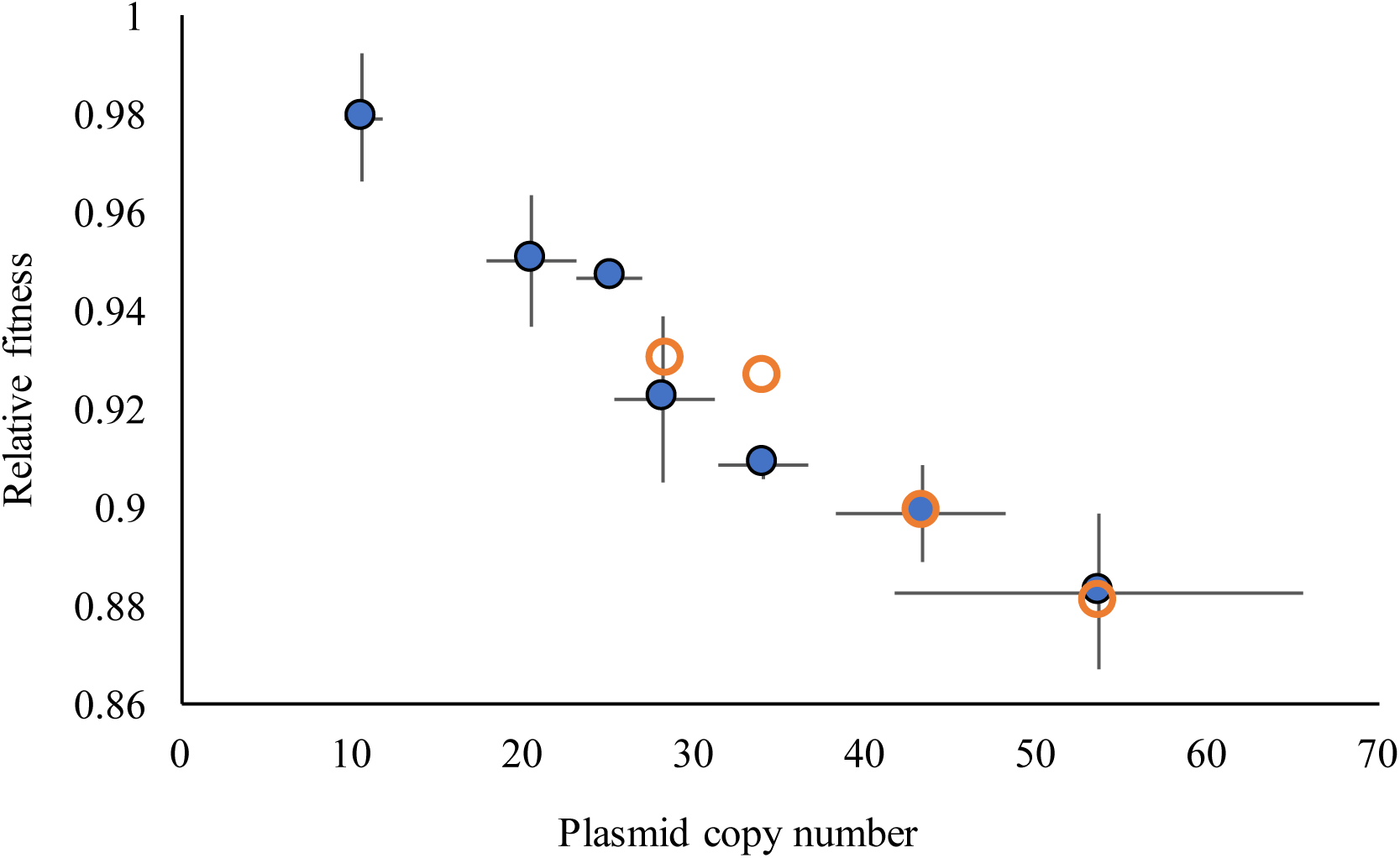
Plasmid copy number and fitness. Correlation between relative fitness of the Rd strain transformed with the different plasmids (Y axis), calculated as 1*+s*, and total plasmid copy number per chromosome (X axis). From left to right the blue circles represent the empirical fitness of: Rd/pB1006, Rd/pB1005, Rd/pB1000, Rd/pB1005/pB1006, Rd/pB1006/pB1000, RdpB1005/pB1000 and Rd/pB1005/pB1006/pB1005. SEM are also indicated. Orange circumferences denote the expected fitness calculated as the multiplication of the fitnesess of the strains bearing plasmids independently. From left to right is represented the theoretical fitness of Rd/pB1005/pB1006, Rd/pB1006/pB1000, RdpB1005/pB1000 and Rd/pB1005/pB 1006/pB 1005.

### The cost of ColE1-like plasmids is proportional to total plasmid copy number

Previous reports showed that the biological cost of a single plasmid, including a ColE1 plasmid (18, 35), is proportional to its copy number in the host cell (19, 36). We measured plasmid copy number (PCN) of pB1000, pB1005 and pB1006 in all the strains using quantitative PCR (qPCR) (Table 1, Figure 3). For all three plasmids, their specific PCN remained equal whether coexisting with the other plasmids or inhabiting the cell alone: pB1000 (ANOVA: P = 0.95; F = 0,10; df = 3, 16), pB1005 (ANOVA: P = 0,79; F = 0,33; df = 3, 16) and pB1006 (ANOVA: P = 0.54; F = 0,75; df =3, 16). These results suggested that replication of these ColE1 plasmids, and therefore PCN control, remained independent despite the high similarity of their conserved region. Interestingly the total PCN, regardless of plasmid type, strongly correlated with the reduction of relative fitness in the host bacteria (Pearson’s test r(19) = 0.90; P < 0.001) (Figure 3). Therefore, the total number of plasmids present in the cell could explained the biological cost imposed by these ColE1 replicons in a non-adapted host, even when the plasmids carried different resistance genes.

**Figure 3.**
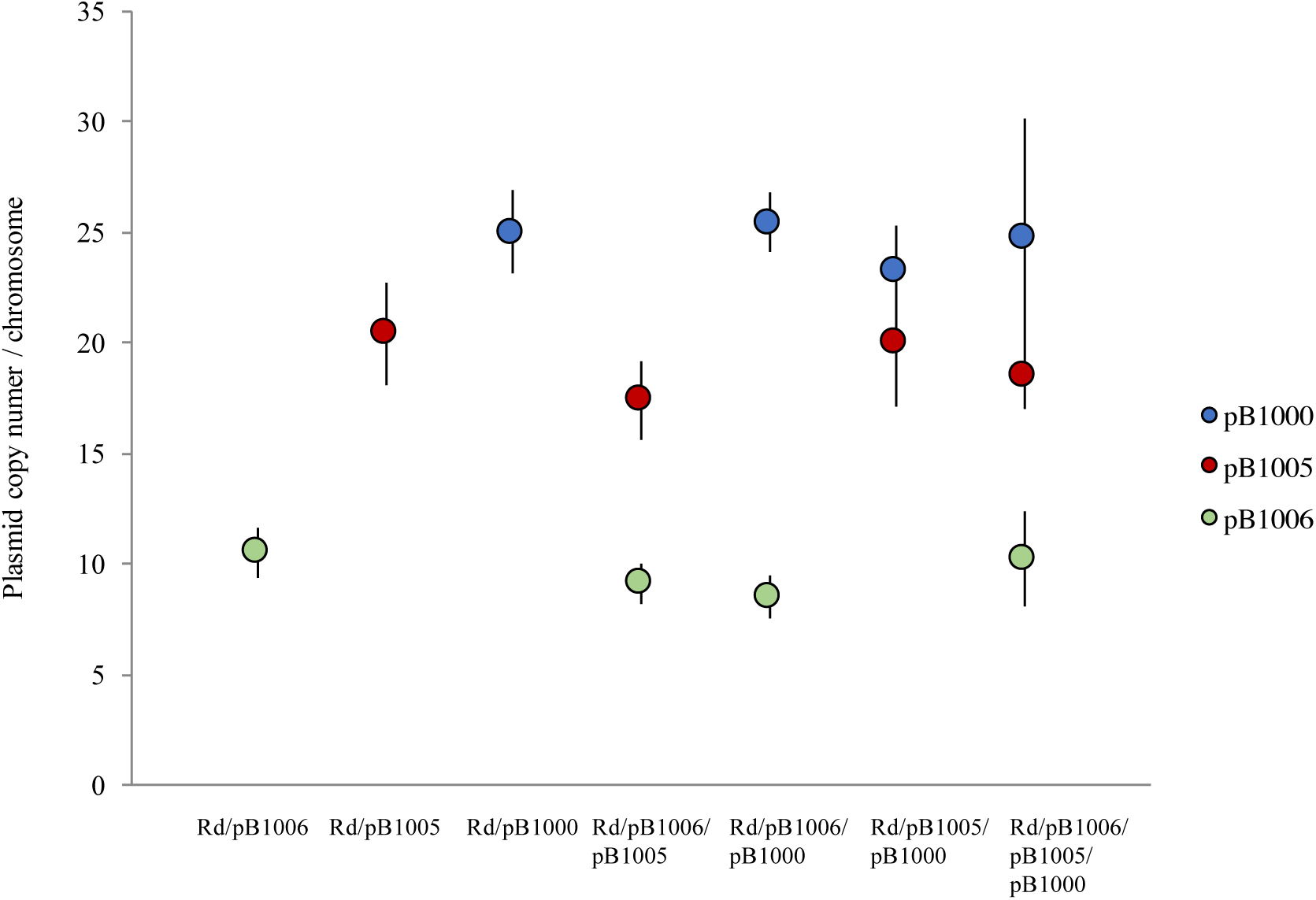
Plasmid copy number in Rd transformed strain. Plasmid copy number per chromosome (Y axis) measured from the seven different combinations (X axis). Green, blue and red dots denote pB1006, pB1005 and pB1000 respectively. SEM are also indicated.

We also analyzed the resistance levels conferred by the plasmids in the different combinations. We measured the minimal inhibitory concentration (MIC) of the three main antibiotics counteracted by the three plasmids (pB1006; tetracycline, pB1005; streptomycin and pB1000; ampicillin). The resistance levels conferred by these plasmids remained constant whether they were alone or coexisting, suggesting that plasmid coexistence did not affect the expression level of these genes (Table 1).

In summary, our results showed that recently acquired ColE1 plasmids acted as independent biological units in the cell, conferring antibiotic resistance, maintaining copy number and imposing fitness costs autonomously.

### Compensatory evolution favors the acquisition of new ColE1 plasmids

Our results suggested that the common coexistence of ColE1 plasmids in nature is not due to positive epistasis alleviating the cost imposed by multiple plasmids. An alternative hypothesis that we propose here is that plasmid coexistence may be promoted by compensatory evolution. The idea underlying this hypothesis is that after the acquisition of a first (ColE1) plasmid by a bacterium, compensatory evolution will eliminate the fitness cost produced this plasmid (22, 37-45). If the cost imposed by different ColE1 plasmids comes from a similar origin, as our previous results suggested, adaptation to this plasmid would facilitate the acquisition of further ColE1 plasmids at no extra cost.

To test this hypothesis, we propagated *H. influenzae* Rd bearing pB1000, which is the plasmid imposing the highest fitness cost (approximately 5% reduction in relative fitness), for 200 generations. Rd/pB1000 increased 18% its relative fitness after 100 and 25% after 200 generations. We selected 1 clone from the Rd/pB1000 population after 100 and 200 generations and we transformed pB1006 and pB1005 separately and together in these clones. We measured the relative fitness of all these clones and, interestingly, there were no significant differences among the relative fitnesses of Rd/pB1000 clones and the new clones bearing also pB1005, pB1006 and pB1005/pB1006 at generation 100 (ANOVA P = 0,85 F = 0,16 df = 2,6) or 200 (ANOVA P = 0,92 F = 0,16 df = 2,6) (Figure 4). In addition, there were no differences in the plasmid copy number across the experiment: pB1000 (ANOVA P = 0.75, F = 0.29 df = 2,12), pB1005 (ANOVA P = 0.25, F = 1.83 df = 2,5) and pB1006 (ANOVA P = 0.71, F = 0.36 df = 2,6) showing that the reduction of the fitness cost of the replicons were not related with a decrease in the plasmid copy number.

**Figure 4.**
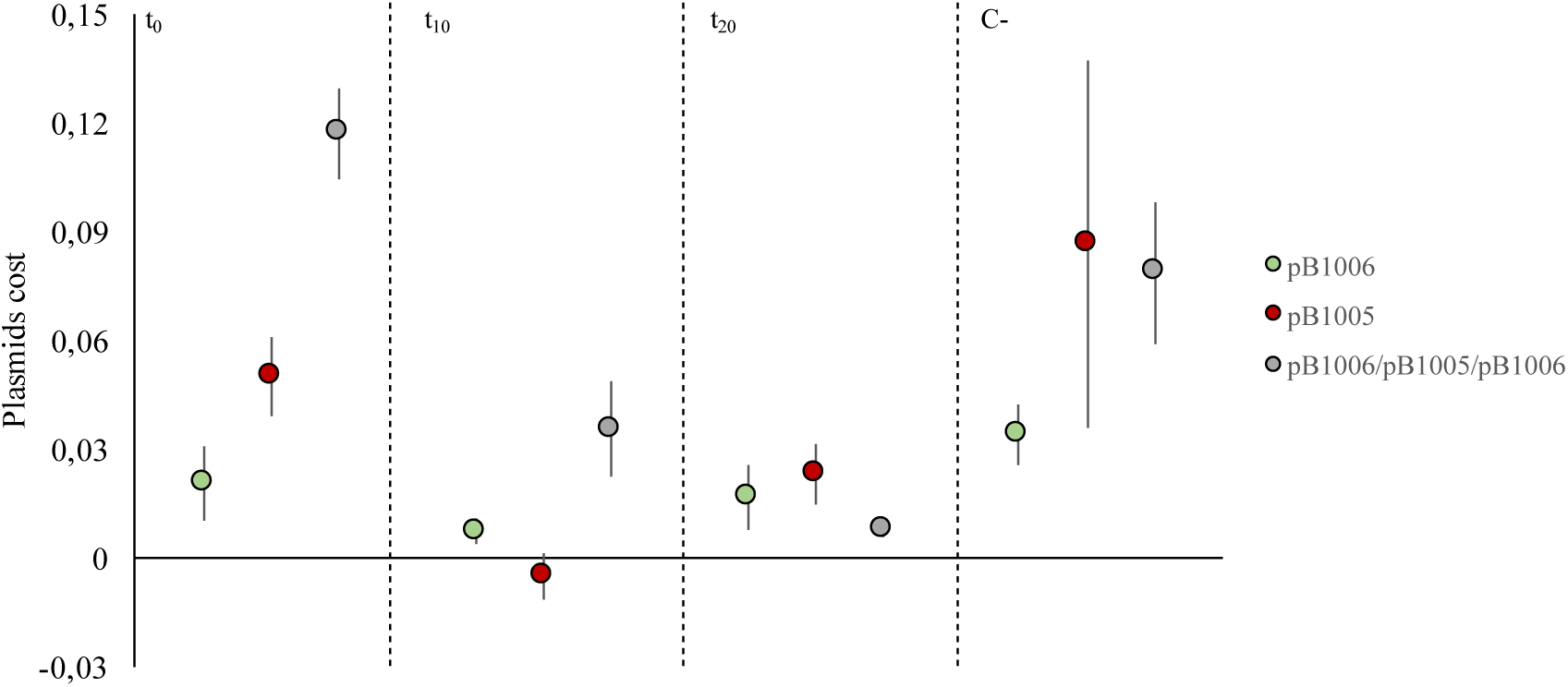
Plasmids cost across the experiment Rd/pBl000. Fitness cost (±SEM) of pB 1006 (green), pB1005 (red) and pB1000/pB1005/pB1000 (grey) calculated as the selection coefficient (see methods) at different time-points of the experiment. In to, the dots represent the fitness cost of the plasmids when newly acquired by Rd strain. In t_100_ and t_200_ is represented the biological cost of the plasmids when transformed in the Rd/pB1000 evolved during 100 and 200 generations. In C-is represented the biological cost of the plasmids transformed in the Rd evolved without pB1000 during 100 generations.

To confirm that the absence of costs associated to additional plasmid carriage in our experiment was a result of compensatory evolution and not only a result of general adaptation to the experimental conditions, we propagated the Rd strain lacking the plasmid in the same conditions as described above during 100 generations. This evolved Rd strain increased its relative fitness approximately 21.5% (SEM = 0,011) compared to the ancestral Rd (Figure 4). We selected a clone from the population and we electroporated pB1006, pB1005, pB1000 and pB1006/pB1005/pB1000. The fitness cost (*s*) of the plasmids in the resulting strains were: pB1006 *s* = −0,034 (SEM ± 0.008), pB1005 *s* = −0.086 (SEM ± 0.050), pB1000 *s* = −0,045 (SEM ± 0.018) and pB1006/pB1005/pB1006 *s* = −0,078 (SEM ± 0.019). Therefore, the plasmids imposed the same biological cost as in the non-evolved strain: pB1006 (two-tailed t-test P = 0.55; t = −1.92; df = 4), pB1005 (two-tailed t-test P = 0.57; t = −0.97; df = 4), pB1000 (two-tailed t-test P = 0.62; t = −0.80; df = 4) and pB1006/pB1005/pB1000 (two-tailed t-test P = 0.29; t = −0,40; df = 4).

Taken together, these results showed that compensation of the cost produced by pB1000 facilitated the acquisition of pB1005 and pB1006 cost by the host bacterium at no additional (Figure 5).

**Figure 5.**
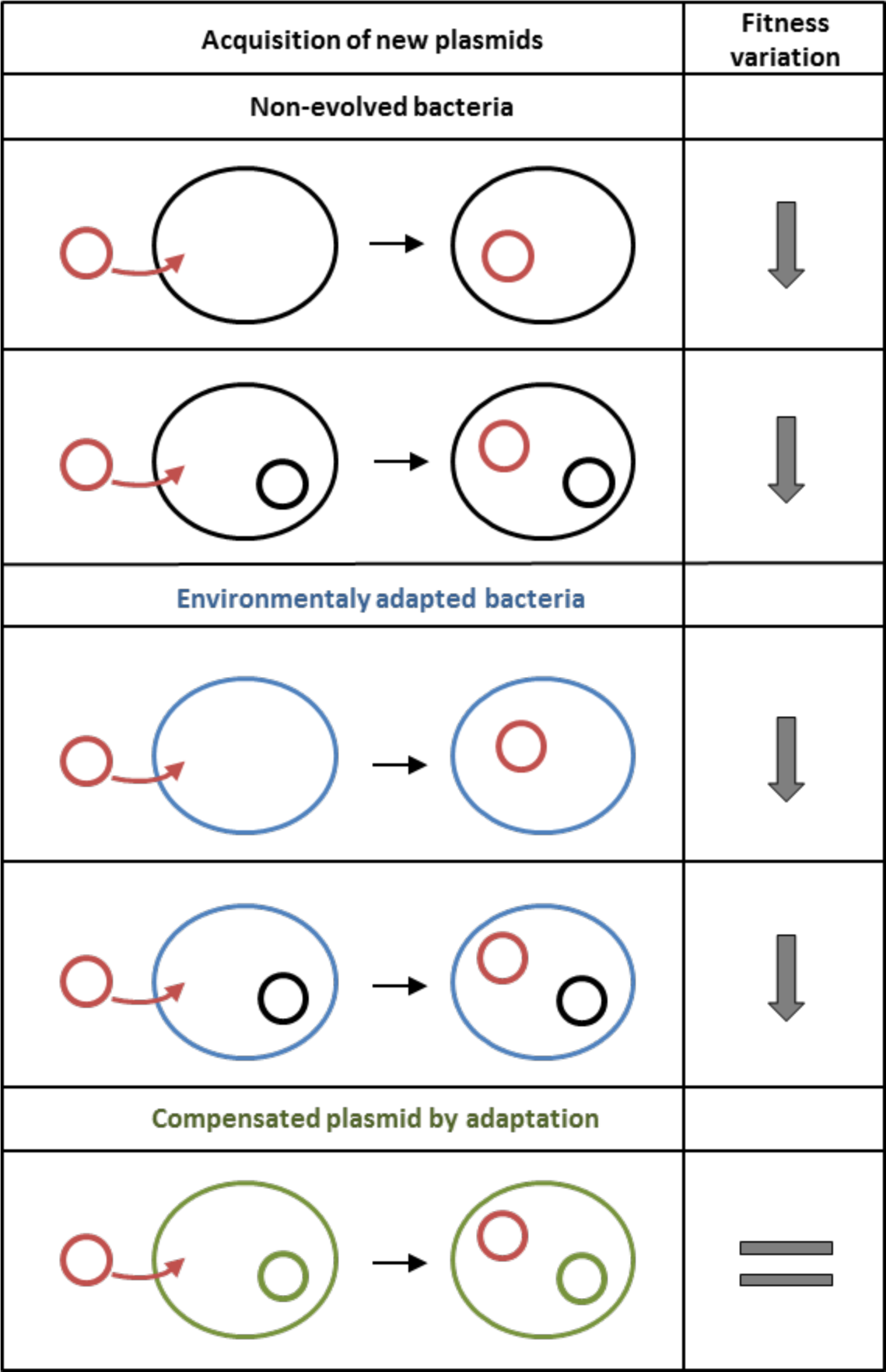
Compensatory evolution facilitates plasmid cohabitation. Acquisition of new plasmid(s) and the effect on the bacterial relative fitness. Red circle denote the acquired plasmid. Non-evolved bacteria or non-evolved plasmid are shown in black; laboratory adapted bacteria are shown in blue and compensated bacteria-plasmid are shown in green. The acquisition of a new ColE1 plasmid in the naïve host (bearing or lacking a plasmid), entails a decrease in the bacterial relative fitness, as well as the acquisition of the plasmid in an environmental-adapted bacterium lacking a plasmid. The acquisition of a new plasmid in the compensated plasmid-carrying bacteria does not result in any fitness cost.

## Conclusion

This study provides new information regarding the biology, cost, persistence and maintenance of multicopy plasmids in bacterial populations. ColE1 plasmids tend to coexist in natural bacteria, conferring antibiotic multiresistance (9, 13, 14). Our results suggest that compensatory evolution mitigating the initial cost produced by a ColE1 plasmid pre-adapts the host bacterium to acquire extra ColE1 plasmids. Although we have only explored the case of plasmids ColE1, a recent report showed that mutations compensating for a particular plasmid also alleviated the cost of plasmids from different families (23). Therefore, we argue that the ability of different bacteria to compensate the costs produced by plasmids may play a key role shaping the distribution of plasmids across bacteria (46).

## Acknowledgment

This work was supported by REDEEX-2 (MICINN, BFU 2011-14145-E) and EU projects EvoTAR 282004-FP7 and EFFORT 613754-FP7. We thank Natalia Montero for excellent technical assistance.

## Author contribution

AS-L, ASM and BG-Z conceived and designed the experiments; AS-L, MA-A, CB-B performed the experiments; AS-L, ASM, and BG-Z analyzed the data; RO-H contributed reagents/materials analysis tools; AS-L, AH, ASM and BG-Z wrote the paper; BG-Z approved the final version of the manuscript.

---

**Figure S1.**
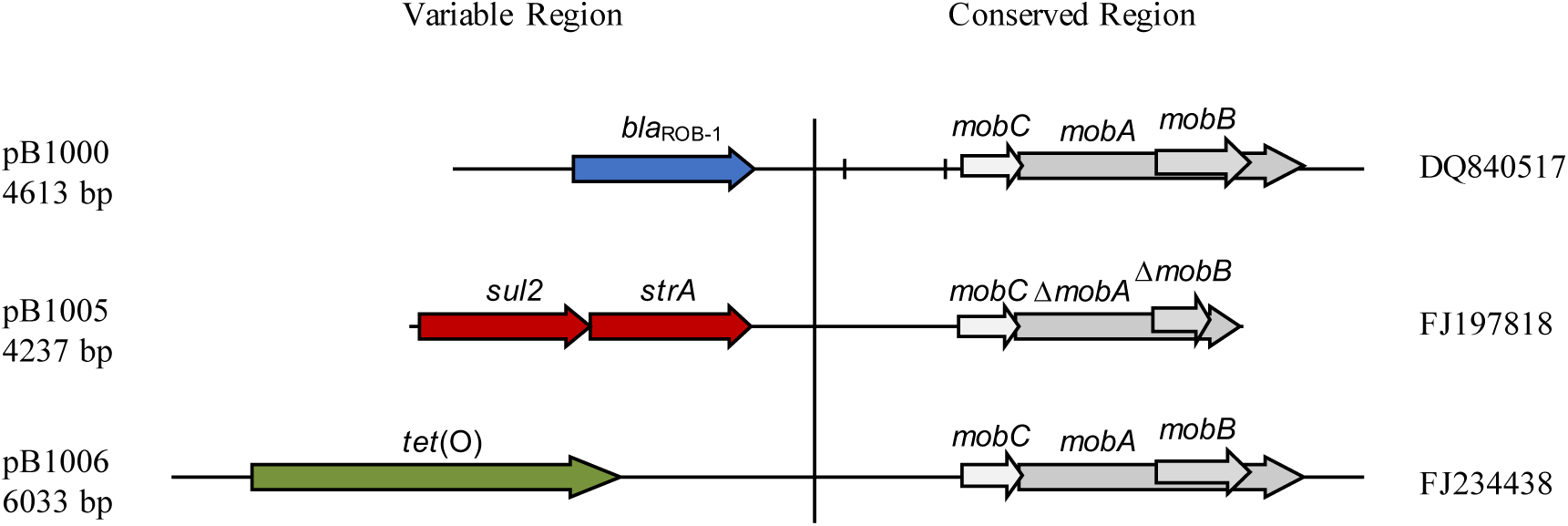
Genetic structure of ColE1 plasmids utilized in this study. Schematic diagram of the 3 ColE1 plasmids used in this study. The reading frames for genes are shown as arrows, with the direction of transcription indicated by the arrowhead. The names of the genes are indicated. Genes encoding plasmid relaxases are shown in gray and the *rom* gene implicated in the regulation of plasmid replication is shown in yellow. In pB1000, two vertical bars bracket the region containing the origin of replication *(oriV)*. The large vertical bar separates the conserved region of the plasmids, to the right, from the variable region of the plasmids, to the left. The accession numbers of plasmids are also indicated. The origins of replication *(oriVs)* of these three plasmids are highly similar compared to the unique *oriV* described in a ColEl from Pasteurellaceae family (28): 89.43%, 99.93% and 91.01% conservation in pB1000, pB1005 and pB1006 respectively.

**Table S1.**
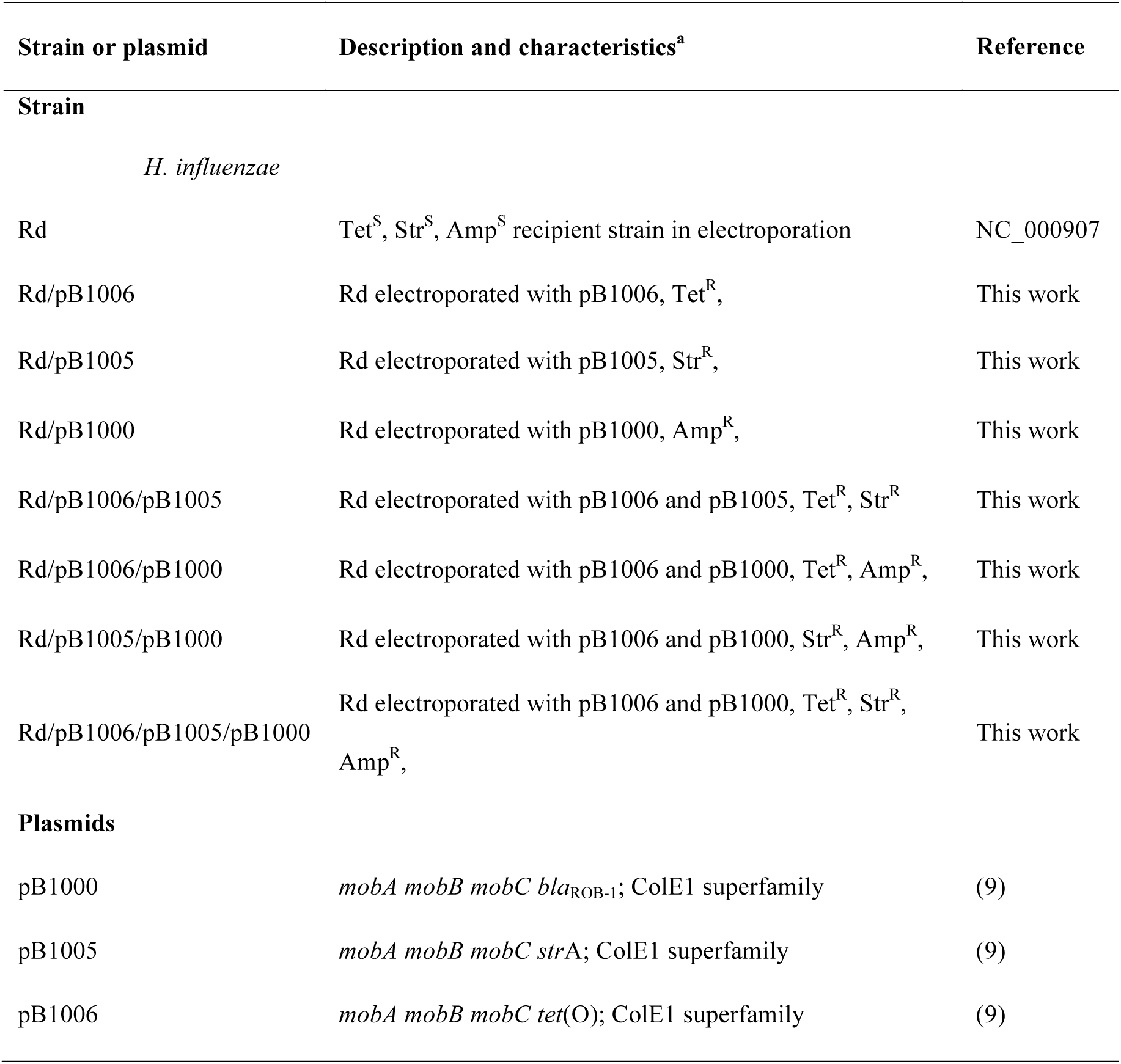
Strains and plasmids used in this study.

Strains and plasmids used in the compensation-acquisition hypothesis. ^a^ Tet, tetracycline; Str, streptomycin and Amp, ampicillin; “R” and “S” means resistant and susceptible respectively.

**Table S2.**
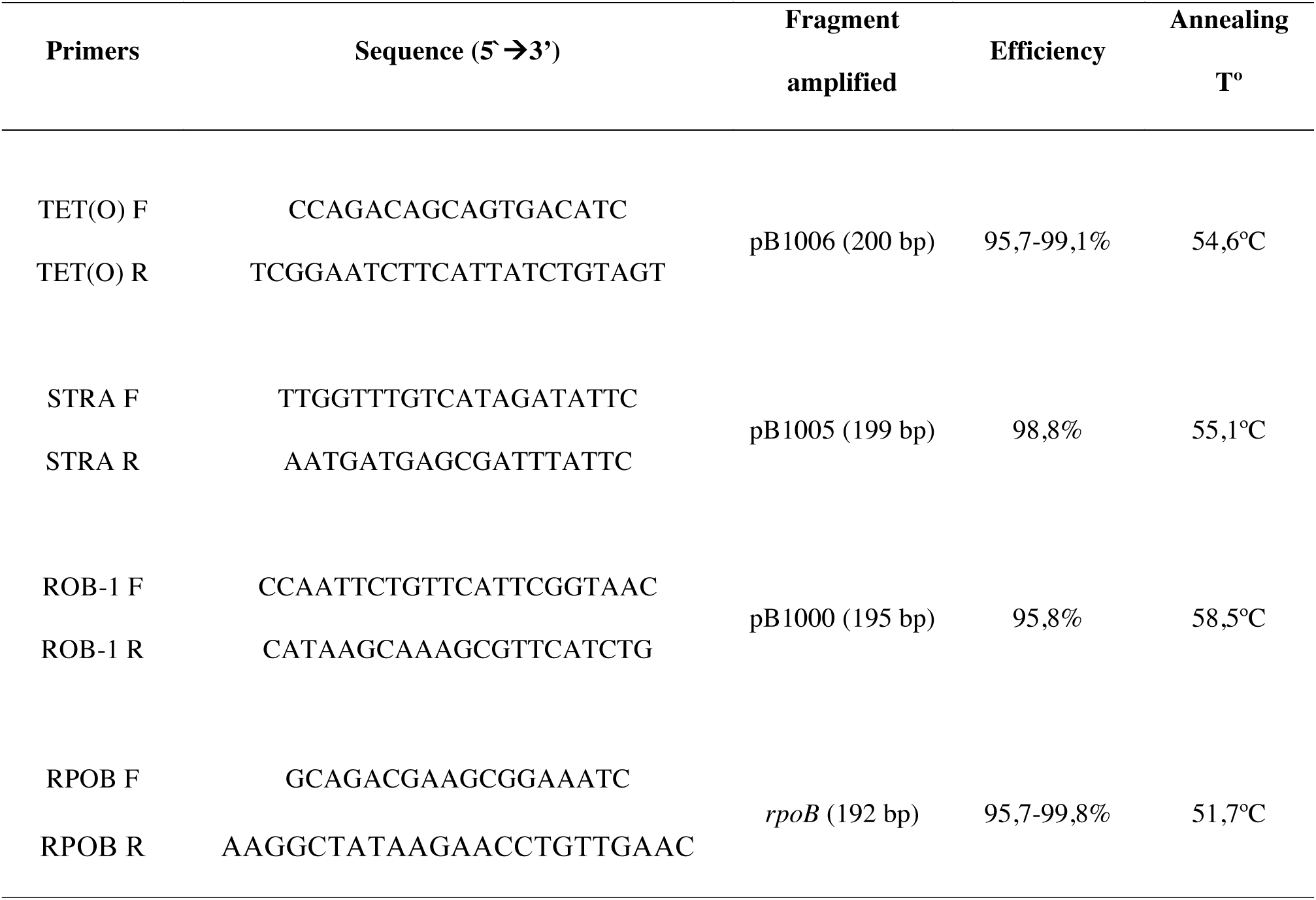
Primers used in this study.

Primers used in the qPCRs. The sequence, fragment amplified, efficiency of the qPCR and Annealing T° of the primers are indicated.

**Table S3.** ColE1 plasmids cohabiting in nature.

| Specie | Strain | ColE1-like plasmid | Others<sup>a</sup> | Characteristics<sup>b</sup> |
| --- | --- | --- | --- | --- |
| **Enterobacteriaceae** |  |  |  |  |
| *Citrobacter freundii* | CAV1741 | pCAV1741-1960,<br>pCAV1741-3233 | 4 | HP<sup>1</sup>, HP |
| *C. freundii* | CAV1321 | pCAV1321-1916,<br>pCAV1321-3233,<br>pCAV1321-3820,<br>pCAV1321-4310,<br>pCAV1321-4938 | 4 | HP, HP, HP, HP,<br>HP |
| *Enterobacter cloacae* | 34978 | P34978-4.398, p34978-5.413, p34978-2.725 | 3 | HP, HP, HP, HP |
| *Escherichia coli* | O26:H11 | pO26_4, pO26_3 | 2 | HP, R/M<sup>2</sup> |
| *E. coli* | SMS-3-5 | pSMS35-8, pSMS35_3 | 2 | Col-E10<sup>3</sup>, IS1 |
| *E. coli* | O55:H7 | p12579_5, p12579_4,<br>p12579_3 | 2 | M/R; Aph(3'')-ib,<br>Aph(6)-Id; TEM-1 |
| *E. coli* | E24377A | pETEC_6, pETEC_5 | 4 | Aph(6)-Id,<br>Aph(3'')-Ib, *sul2*;<br>*nga* |
| *E. coli* | SE11 | pSE11-5, pSE11-4 | 4 | HP, *nga* |
| *E. coli* | PCN061 | PCN061p3, PCN061p2,<br>PCN061p1 | 3 | *sul2*, Aph(3'')-Ib,<br>Aph(6)-Id; HP; HP |
| *E. coli* | ST648 | pEC648_7, pEC648_5 | 4 | HP, HP. |
| *Klebsiella oxytoca* | CAV1374 | pCAV1374-6538,<br>pCAV1374-14 | 9 | Tn3, Tn3 |
| *K. pneumoniae* | 234-12 | pKpn2312-5, pKpn23412-4 | 0 | HP, HP |
| *K. pneumoniae* | MGH 78578 | pKPN6, pKPN7 | 3 | HP, HP |
| *K. pneumoniae* | Kp13 | pKP13b, pKP13c | 4 | SMR<sup>4</sup> |
| *Salmonella*<br>*Typhimurium* | CFSAN001921 | unnamed 2 (NC_021843),<br>unnamed3 (NC_021816) | 1 | HP, HP |
| *S. enterica* | YU39 | pYU39_4.8, pYU39_4.2 | 4 | HP, HP |
| *Serratia marcescens* | CAV1492 | pCAV1492-3223,<br>pCAV1492-6393 | 4 | ND |
| *Shigella flexneri* | 2002017 | pSFxv_3, pSFxv_4,<br>pSFxv_5 | 2 | *strA*, HP, Fe<sup>5</sup> |
| *S. sonnei* | Ss046 | pSS046_spA, pSS046_spB | 2 | *tet(A)*, Col-E1<sup>3</sup> |
| **Pasteurellaceae** |  |  |  |  |
| *A. pleuropneuminae* | AP76 | pB1002, pB1003, APP7_C | ND | *bla*<sub>ROB-1</sub>; *strA*, *sul2*; * |
| *H. parasuis* | BB1020 | pB1000, pB1006 | ND | *bla*<sub>ROB-1</sub>, *tet(O)* |
| *P. aerogenes* | BB1084 | pB1000, pIG1 | ND | *bla*<sub>ROB-1</sub>, *strA* |
| *P. multocida* | BB1038 | pB1000, pB1005 | ND | *bla*<sub>ROB-1</sub>; *strA*, *sul2* |
| *P. multocida* | BB1039 | pB1000, p9956 | ND | *bla*<sub>ROB-1</sub>, *tet(H)* |
| *P. multocida* | BB1046 | pB1002, pB1003 | ND | *bla*<sub>ROB-1</sub>; *strA*, *sul2* |
| *P. multocida* | BB1044 | pB1000, pB1005, pB1006 | ND | *bla*<sub>ROB-1</sub>; *strA*, *sul2*;<br>*tet(O)* |

^a^ Non ColE1 plasmids presented in the same cell

^b^ Genes founded in the variable región of ColE1 plasmids: ^1^ HP, Hypothetical Protein; ^2^

M/R, Restriction/Modification System; ^3^ Col-E1, Colicin E1; ^4^ SMR, *Small Multidrug Resistance proteíns;* ^5^ Fe, involved in ferric resistance; ^*^ Cryptic plasmid.

